# SPIAT: An R package for the Spatial Image Analysis of Cells in Tissues

**DOI:** 10.1101/2020.05.28.122614

**Authors:** Tianpei Yang, Volkan Ozcoban, Anu Pasam, Nikolce Kocovski, Angela Pizzolla, Yu-Kuan Huang, Greg Bass, Simon P. Keam, Paul J. Neeson, Shahneen K. Sandhu, David L. Goode, Anna S. Trigos

## Abstract

Spatial technologies that query the location of cells in tissues at single-cell resolution are gaining popularity and are likely to become commonplace. The resulting data includes the X, Y coordinates of millions of cells, cell phenotypes and marker or gene expression levels. However, to date, the tools for the analysis of this data are largely underdeveloped, making us severely underpowered in our ability to extract quantifiable information. We have developed SPIAT (**Sp**atial **I**mage **A**nalysis of **T**issues), an R package with a suite of data processing, quality control, visualization, data handling and data analysis tools. SPIAT includes our novel algorithms for the identification of cell clusters, cell margins and cell gradients, the calculation of neighbourhood proportions, and algorithms for the prediction of cell phenotypes. SPIAT also includes speedy implementations of the calculation of cell distances and detection of cell communities. This version of SPIAT is directly compatible with Opal multiplex immunohistochemistry images analysed through the HALO and InForm analysis software, but its intuitive implementation allows use with a diversity of platforms. We expect SPIAT to become a user-friendly and speedy go-to package for the spatial analysis of cells in tissues.

SPIAT is available on Github: https://github.com/cancer-evolution/SPIAT

## Introduction

Recent technological advances in spatial technologies in the last 2-3 years, such as in multiplex immunohistochemistry, microscopy and spatial transcriptomics, provide detailed spatial information at single-cell resolution, moving the field into the quantitative arena. A popular application of these spatial technologies has been in the study of the tumour microenvironment. Immune cells in the tumour area have been found to display distinct distribution patterns linked with survival in a number of solid tumours (1, 2). For example, control of tumour growth by the immune system is linked to high levels of lymphocytes in close proximity to tumour cells (immune infiltration), whereas immune cells in the periphery of the tumour are considered to be excluded. The first profile has been linked to better prognosis (3), and studies have shown a link between response to immune checkpoint inhibitors and high levels of lymphocytes in the tumour invasive margin and tertiary lymphoid structures (4-7). This has resulted in a great appetite for the study of the spatial patterns of cells in the microenvironment, even beyond immune cells.

The new Opal™ multiplex immunohistochemistry staining protocol allows the use of 6-8 markers simultaneously on a single slide followed by imaging on the Perkin Elmer Vectra™quantitative imaging system. It has gained significant popularity due to its applicability to formalin fixed paraffin embedded tissue sections, allowing the examination of samples taken in a clinical setting. The imaging system records the fluorescence emission for each marker, and each cell is assigned an X,Y coordinate of its location in the tissue, effectively providing single-cell resolution. The fluorescence emission maps of individual markers are then combined to yield the location and phenotype of cells, along with marker intensity.

Currently, studies into spatial tissue analysis have been devoted to extracting information from raw microscopy images. Machine learning is commonly used to perform cell segmentation (identifying cells in an image), and tissue segmentation (differentiating stromal and tumour regions in the image). Next, cell phenotyping is performed based on marker intensity (e.g. CD3 for T cells) or a combination of markers (e.g. CD3 and CD4 for helper T cells). It is not atypical to have from a few thousand to several million cells in an image, resulting in large amounts of big data. Unfortunately, once this information has been extracted, the methods for the downstream analysis and interpretation of these data are rudimentary, resulting in their underutilization.

To date, most approaches tackling this issue determine the density of cells within the tumour area or measure the Euclidean distances of all cells against all cells, identifying populations that are closest to tumour cells (8). However, these analytical approaches leave most of the information contained in these images unquantified and unexplored, limiting the questions that can be asked and the reducing the possibility of new discoveries. More advanced studies investigating the patterns of clustering of immune cells revealed a tight link with prognosis, tumour type and response to immunotherapy (1, 2, 9), indicating a largely untapped source for novel biological insights.

We have developed the R package SPIAT for the **Sp**atial **I**mage **A**nalysis of Cells in **T**issues. SPIAT includes a broad range of methods that allow (1) reading in Opal multiplex immunohistochemistry images processed with the two most popular Opal software (InForm™ or HALO™) and quality control, (2) multiple approaches for the visualization of the distribution of cells, either by phenotype, marker intensity levels, and surface plots, (3) metrics for the calculation of mixing of cell phenotypes, (4) identification of cell clusters or communities of cells, (5) identification of cell gradients and (6) cell phenotype predictions.

### Input to SPIAT

## Methods and workflow

An overview diagram of SPIAT is available in Figure 1. Our current version includes over 26 functions. The input to SPIAT are the cell IDs, X, Y coordinates, marker intensities and cell phenotypes (which markers where positive in each cell). Images must have been cell segmented previously, and cells phenotyped using HALO or InForm software, although SPIAT offers the option of calling phenotypes based on marker levels and marker combinations. We have tested SPIAT successfully with images of up to ∼1 million cells, but implementations of the algorithms were made to optimise speed, so there is currently no upper limit. Our input function (*format_image_to_sce*) reads in the format from HALO and InForm software, but these formats are relatively generic, and users can reformat their files to match one of these and then input to SPIAT. Users are also required to input the columns of markers of interest and the location where the marker is expected to be located (nucleus, cytoplasm or membrane).

**Figure 1.**
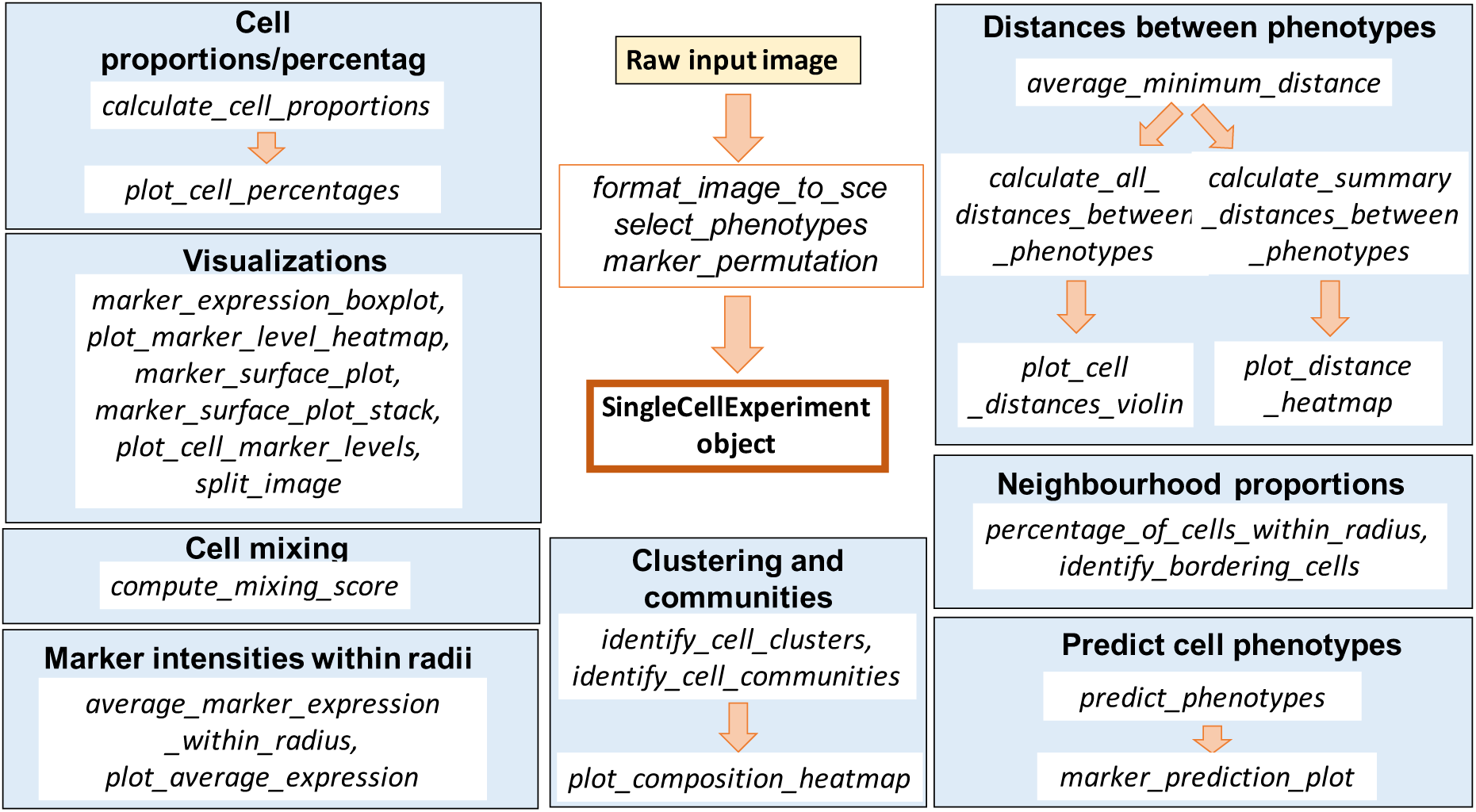
Overview diagram of functions in SPIAT

The base object for SPIAT is SingleCellExperiment, which is designed for single-cell gene expression data. Here the marker intensities are treated as gene expression levels, and the cell coordinates and categorical phenotypes (if available) as additional metadata. For ease of use, we designed the package so most functions are independent of each other, and start from the SingleCellExperiment object (Figure 1). We also implemented *image_splitter*, which allows splitting images into sections defined by the user. This is recommended in the case of large images, or images where there are two independent tissue sections.

### Quality control of images

Since cell phenotypes are defined by combination of markers, low cell segmentation quality, antibody ‘bleeding’ from one cell to another or inadequate marker thresholding, can lead to the assignment of erroneous cell phenotypes. Cells might be identified as being positive for a biologically unfeasible combination of markers (for example, cells positive for a tumour marker and an immune marker, such as CD3). Therefore, SPIAT allows the user to manually input which phenotypes they would like to keep or exclude (*select_phenotypes*).

Alternatively, we also implemented a shuffling strategy to determine how likely it is to obtain a particular combination of markers by chance (*marker_permutation*). Here, we create a null distribution by permuting the marker labels of cells, and calculate the empirical p-value of whether an image is enriched or depleted in a particular combination of markers. This is meant to provide guidance to the users of which combination is likely to occur by chance (e.g. because there are high numbers of cells positive for particular marker), but it is not absolute, and users are highly encouraged to review the results.

We also implemented a function that plots the marker intensities of cells identified as being positive or negative for a given phenotype (*marker_expression_boxplot*). Cells positive for a marker should have higher levels of the marker. Since HALO and InForm use machine learning to determine positive cells, and not a strict threshold, some positive cells will have low marker intensity, and vice versa. However, an unclear separation of marker intensities between positive and negative cells would suggest incorrect phenotyping or unreliable phenotyping due to background noise.

Finally, SPIAT calculates the proportion of cells of each phenotype (*calculate_cell_proportions*), which can be visualized as barplots (*plot_cell_percentages*).

### Visualization of tissues

SPIAT has multiple options for visualization of cells in the tissues:

- *plot_cell_categories*: Dot plots, where each dot corresponds to a cell and cells are coloured by phenotype. One plot summarizes all markers and cell types.
- *plot_marker_level_heatmap*: For large images, there is also the option of ‘blurring’ the image, where the image is split into multiple small areas, and marker intensities are averaged within each.
- *plot_cell_marker_levels*: Here the intensity of each marker can be visualized individually, which can be used to check for an uneven staining or high background intensity.
- *marker_surface_plot*: Plots the level of markers as a 3D surface plot. To stack the surface plots of multiple markers in a single plot we have implemented *marker_surface_plot_stack.* This allows the identification of co-occurring and mutually exclusive markers.

### Distances between cell types

SPIAT implements a number of functions for the calculation of Euclidean distances between cells. First, we can compare the locations of two phenotypes (phenotype A and phenotype B) by identifying the closest cell of phenotype B to each of the cells of phenotype A. This creates a distribution of minimum distances between phenotypes A and B (*calculate_all_distances_between_phenotypes*), which can be visualized as a violin plot (*plot_cell_distances_violin*). Visualization of this distribution often reveals whether pairs of cells are evenly spaced across the image, or whether there are clusters of pairs of phenotypes. As a summary statistic, SPIAT also calculates the mean, median and standard deviation of each combination of phenotypes (*calculate_summary_distances_between_phenotypes*), with visualization of cell distances as a heatmap (*plot_distance_heatmap*).

### Detections of cell clusters and communities

One of the main recurrent questions in the analysis of the spatial distribution of cells in the microenvironment is the presence or absence of aggregates of a particular phenotype or combination of phenotype. In SPIAT, we make the distinction of two types of cell aggregates. The first are “clusters”, which are cell aggregates composed of non-tumour cells, often by immune cells. For the detection of clusters, *identify_cell_clusters* only considers cell phenotypes of interest defined by the user (e.g. clusters of CD4+ cells). Euclidean distances between cells are calculated, and pairs of cells with a distance less than a threshold are considered to be ‘interacting’, with the rest being ‘non-interacting’. Hierarchical clustering is then used to separate the clusters.

While we recommend users to test out different thresholds and then visualise the clustering results, we also offer the *average_minimum_distance* function, which calculates the average minimum distance between all cells in an image to use as a reference or starting point.

The second type of cell aggregates are “communities”, as previously defined (9). Here, communities correspond to micro-niches or micro-ecosystems of cells that are geographically located close to each other (*identify_cell_communities*). The main distinction between clusters and communities is that the algorithm for the detection of communities does not take into account cell phenotype. Therefore, communities often consist of a combination of different cell types, including tumour cells. PhenoGraph (10) is used as the clustering algorithm to detect communities.

SPIAT also includes the *plot_composition_heatmap* function, which allows visualization of the cell composition of clusters or communities. This allows discerning whether there are regions of heterotypic cell-cell interactions, or whether the image is highly structured with poor cell mixing.

### Mixing scores

SPIAT includes calculation of cell mixing scores, which was originally defined as the number of immune-tumor interactions divided by the number of immune-immune interactions (2). We have generalized this score to allow calculation of any two cell phenotypes defined by the user.

### Identification bordering cells

A common question that arises in the study of the tumour microenvironment is whether immune cells are close to the tumour margin, or how does the density of a particular cell type differ between the tumour and stromal areas. A first step in answering this question is the identification of cells that separate two distinct tissue areas (e.g. tumour area vs. stromal area). This tissue segmentation process often relies on machine learning or manual delineation of the borders in HALO or InForm, which are either coarse and/or time-consuming, rely on user intervention and often result in poor resolution. In SPIAT we developed an algorithm (*identify_bordering_cells*) to automatically detect bordering cells, with only 5 parameters: *reference_marker* (reference area to be identified, i.e. tumour marker), *noise_radius, radius, lower_bound* and *upper_bound*. The algorithm is as follows:

1. <u>We select</u>*<u>reference_marker</u>* <u>cells surrounded by other</u>*<u>reference_marker</u>* <u>cells:</u> We first identify *reference_marker* cells with other *reference_marker* cells within *noise_radius*.
2. <u>We select non-reference cells surrounded by other non-reference cells:</u> Only keep non-reference cells with non-reference cells within *noise_radius*.
3. <u>We identify bordering cells as those tumour cells surrounded by a certain percentage of stromal cells:</u> We combine the cells identified in (1) and (2) and identify all neighbouring cells within *radius.* We then calculate the percentage of stromal cells in the neighbourhood of each tumour cell, and mark those tumour cells with a percentage of stromal cells within the *lower_bound* and *upper_bound* as bordering cells.

The identified bordering cells can then be used as a reference for calculation of distances to other cell types. Note that while here we use tumour cells as an example for reference cell, the same can be applied to any cell type.

### Identifying gradients of cells

One of the main questions in the spatial analysis of cells in tissues is whether a particular cell type is close to or interacting with another, and this is used as a benchmark to compare groups of images (e.g. primary vs. metastatic). For example, in the study of recognition of tumour cells by the immune system, a main question is whether cytotoxic T cells are close to tumour cells, or whether PDL1+ cells are aggregated in the tumour area.

A common solution is the calculation of the average minimum distance of tumour cells to immune cells, or vice versa. In this case, the minimum distance between an immune cell and a tumour cell is calculated, and then the processed is repeated for all immune cells. This result is then averaged, or the distributions are compared between images using a Wilcoxon test or similar. A major drawback of this naïve approach is that the total number of cells is not considered – even with no preferential attraction of immune cells to tumour cells, a large number of cells will result in smaller minimum distances, resulting in false positive results. In summary, these naïve metrics do not take into account the actual spatial distribution of cells.

In SPIAT we address this challenge using the concept of gradients. If there is a true aggregation or attraction of cell phenotype A to cell phenotype B, then we would see higher levels of marker A closer to cells of phenotype B, with this value decreasing the further we move away from cells of phenotype B. This concept of gradients also allows the identification of repulsion between cell types, such that if we see that marker A intensities are lower when close to cells of phenotype B, but increase as we move away, then we can say that there is likely to be a process of repulsion. This concept is implemented in SPIAT in the *plot_average_expression* function, with the *average_marker_expression_within_radius* being the helper function for the calculation of the marker level within a specified radius.

### Calculation of neighbourhood proportions

SPIAT also offers an additional alternative to characterising cell aggregation, which is the calculation of the neighbourhood proportion (*percentage_of_cells_within_radius*). Here, we define the proportion of a target cell type within the neighbourhood of a reference cell type within a defined radius. This algorithm is inspired by previous work (8), but in that case the average proportion of all cells in an image was used. Here, we perform the calculation for each cell, and thus we take into account differences in spatial distribution. With our method, spatial structures can be identified by pinpointing cells with high neighbourhood proportions of the target cell type.

Of note, *percentage_of_cells_within_radius* can be used for the detection of gradients instead of *average_marker_expression_within_radius*, however we recommend using the latter given its implementation allows a speedier analysis, and it does not depend on cell phenotyping as it uses marker intensities.

### Prediction of phenotypes based on marker levels

One of the main applications of InForm and HALO is for cell phenotyping. Here, the user selects ∼10 cells that are visually ‘positive’ for a particular marker, which are then used as a training set to phenotype the rest to the cells. However, this is usually an iterative process, whereby the user often needs to recalibrate the model by further selecting more cells and so forth. As a result, cell phenotyping is often time-consuming. In SPIAT we have implemented algorithms for the automatic phenotyping of cells based on marker intensities (*predict_phenotypes*) that do not require user intervention or the manual setting of thresholds.

Conceptually, our base algorithm assumes that most cells in an image are not positive for the particular marker of interest. With this assumption, we can estimate the background levels of the marker based on the distribution of marker intensities. We have observed that in most cases, marker levels follow a Beta (or Beta-like) distribution skewed to the left with a long right tail. With the assumption that most cells are negative for the marker, cells in this right tail are marked as being positive. The cutoff is selected as the inflection point of the distribution as it flattens. We have also accounted for cases where there is a weak antibody signal, and the distribution might resemble a normal distribution, or where there is a bimodal distribution, and only cells after the second peak should be considered.

In cases where the cells to be phenotyped are likely to represent most of the cells in the image, for example tumour cells in a tumour-dense tissue, we have added an additional step to our base algorithm. Here, we first phenotype cells based on non-tumour markers (*baseline_markers*) (for example, immune markers). This population of cells is used to determine the distribution of background levels of the tumour marker. Subsequently, we select the 0.95 quantile of the tumour marker in this population as a putative threshold (threshold 1). We empirically determine the inflection point of the distribution of tumour markers using our base algorithm (threshold 2). Finally, we use whichever of the thresholds is greater as a cutoff for positive and negative cells.

SPIAT includes the *marker_prediction_plot* function, which plots the predicted cell phenotypes and the ones obtained using HALO or InForm, for comparison. Of note, this algorithm does not take into account cell shape or size, so if these are required for phenotyping, manual phenotyping with HALO or InForm is encouraged.

## Results

To showcase the power of SPIAT in characterising the spatial distribution of cells in tissues, we include results from 4 test images generated with Opal multiplex immunohistochemistry (Figure 2). One corresponds to a primary melanoma image analysed on the HALO system (Figure 2a), while the others correspond to prostate cancer images with distinct pathologies analysed on InForm: one with highly glandular composition of tumour regions (Figure 2b), another with more diffused distribution (Figure 2c), and one with a clear tumour margin and little immune infiltration (Figure 2d). We used the SOX10 and AMACR markers for melanoma and prostate cancer cells, respectively. Immune markers were DAPI, CD3, CD4, CD8, PDL1 and CD103 for the melanoma image, and DAPI, CD3, CD4, CD8, FOXP3 and PDL1 for the prostate cancer images.

**Figure 2.**
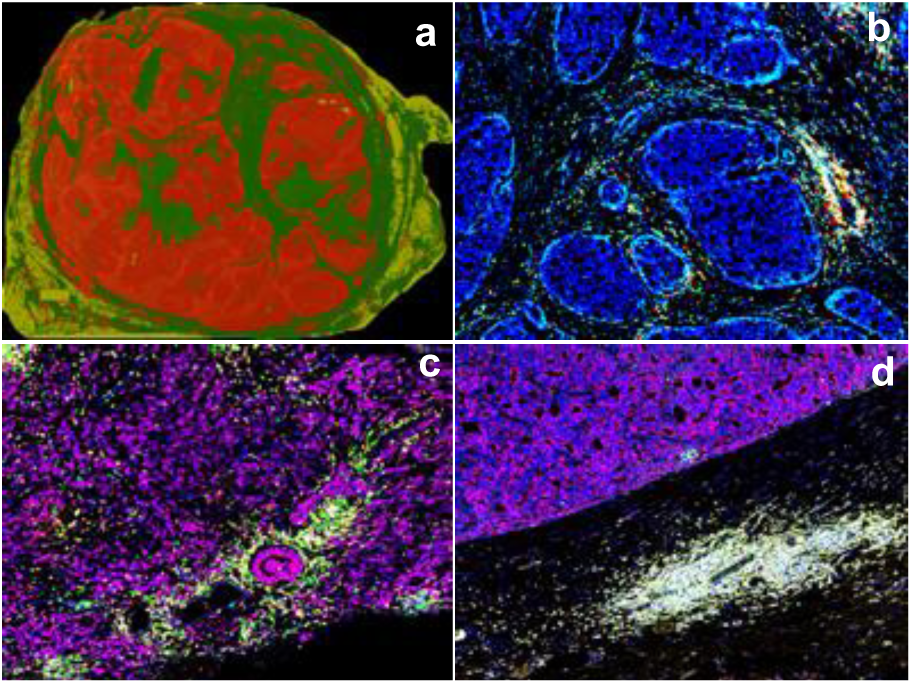
Opal multiplex immunohistochemistry images from (a) primary melanoma, (b-d) primary prostate cancer. In (a), red corresponds to the tumour area. The prostate tumor marker, AMACR, is shown in blue in (b), and purple in (c) and (d). Other colours correspond to immune cells.

After reading in the images and converting them to the single-cell experiment object, we next perfomed quality control of the images. We compared the intensity of each marker in cells phenotyped as positive or negative for the CD4 marker. Cells positively phenotyped were found to have higher CD4 marker intensities (Figure 3a), consistent with an adequate phenotyping. Note that some negative cells had high marker intensities. This is because the phenotyping method used by InForm and HALO do not solely use marker intensities for thresholding, but also cell shape, nucleus size, etc. However, the general trend of higher marker intensities in positive cells should be fulfilled.

**Figure 3.**
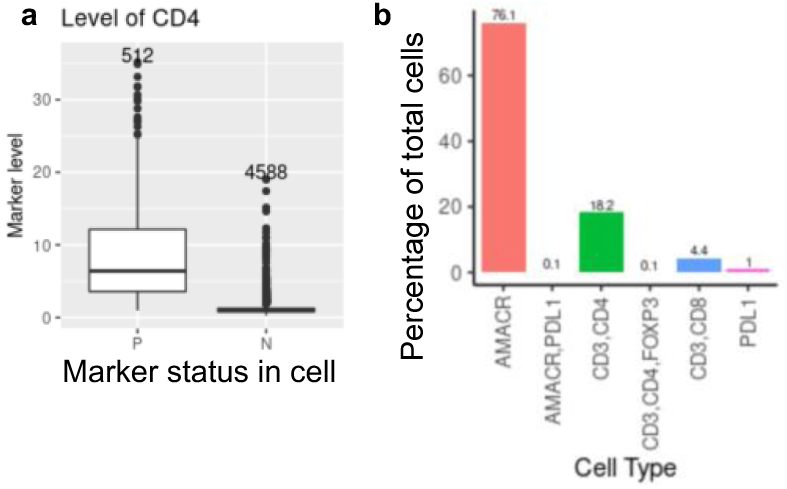
Basic quality control metrics of images. (a) Boxplots of the CD4 marker level of the prostate cancer tissue shown in Figure 2b. Cells positive or negative for a marker were defined in InForm. Cells marked as positive for a marker should have higher intensity levels for that marker, as shown here for CD4. (b) Percentage of each cell phenotype in the prostate cancer tissue shown in Figure 2b. P=positive. N=Negative.

The calculation of the proportion of cells in images revealed that the vast majority of cells were tumour cells, and with few immune cells, as shown in example Figure 3b. There was a small proportion of cells that appeared to have a biologically impossible combination of markers (e.g. AMACR+ and CD4+), which were excluded from further downstream analyses. While the combination of markers of interest was known *a priori*, random shuffling of markers with *marker_permutation* revealed the implausible combinations as being depleted, flagging them for consideration.

We next made a visual inspection of the images to assess staining quality and its potential implications for cell phenotyping. The three 2D visualization options in SPIAT allowed a quick visual assessment of patterns of immune infiltration and potential confounders (Figure 4). CD4+ and CD8+ cells were more common in the melanoma image (Figure 4a,e,i), although high levels of CD4 background (Figure 4i) might have been a potential confounder in the phenotyping of CD4+ cells. In contrast, the proportion of immune cells is much lower in the prostate images (Figure 4 b-d, f-h, j-l), and staining of the CD4+ was much cleaner. Marker intensities were also be explored using 3D surface plots, as shown in Figure 5. Here, we can clearly see that high levels of SOX10+ (tumour) cells did not co-occur with the CD4+ immune cell marker, but rather were found in between areas of high SOX10 intensity. SPIAT also detected tumour borders, both in clear and challenging images (Figure 6).

**Figure 4.**
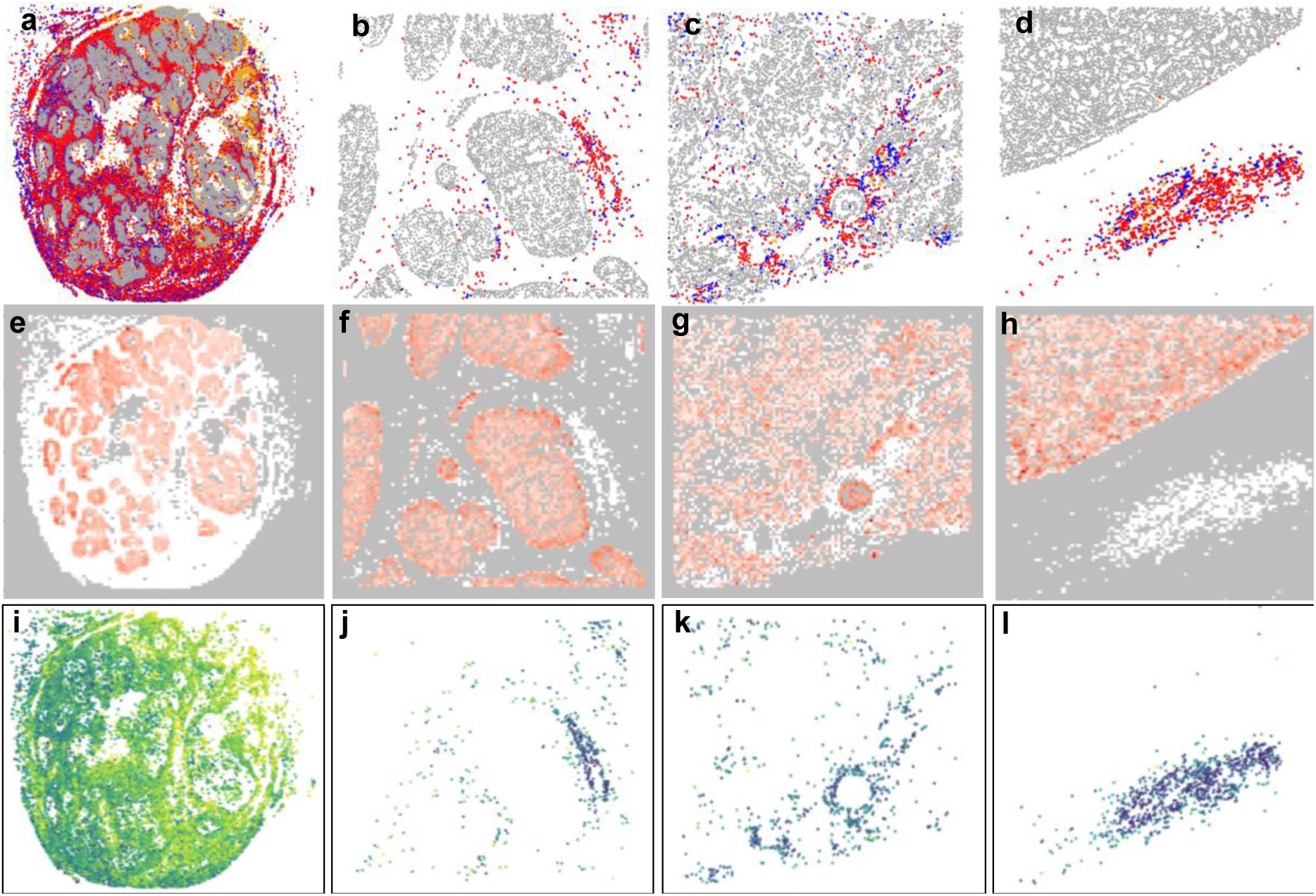
Visualizations available in SPIAT. (a-d) Plot generated with *plot_cell_categories* of each of the melanoma (a) and prostate tissues (b-d). Grey= Tumour cells, Red=CD3+CD4+ cells Blue=CD3+CD8+ cells, Orange=CD103+ cells in the melanoma tissue and PDL1+ cells in prostate cancer tissues. (e-h) Same images as above, but plotting the tumour SOX10 marker in melanoma (e) or the tumour AMACR marker in prostate cancer (f-h) with *plot_marker_level_heatmap.* Red colouring represents the tumour cells, white the immune cells. (i-l) Plots of the CD4 intensities. There are higher background levels of CD4 in the melanoma (i) than in the prostate tissues (j-l). Darker colours indicate stronger marker intensity.

**Figure 5.**
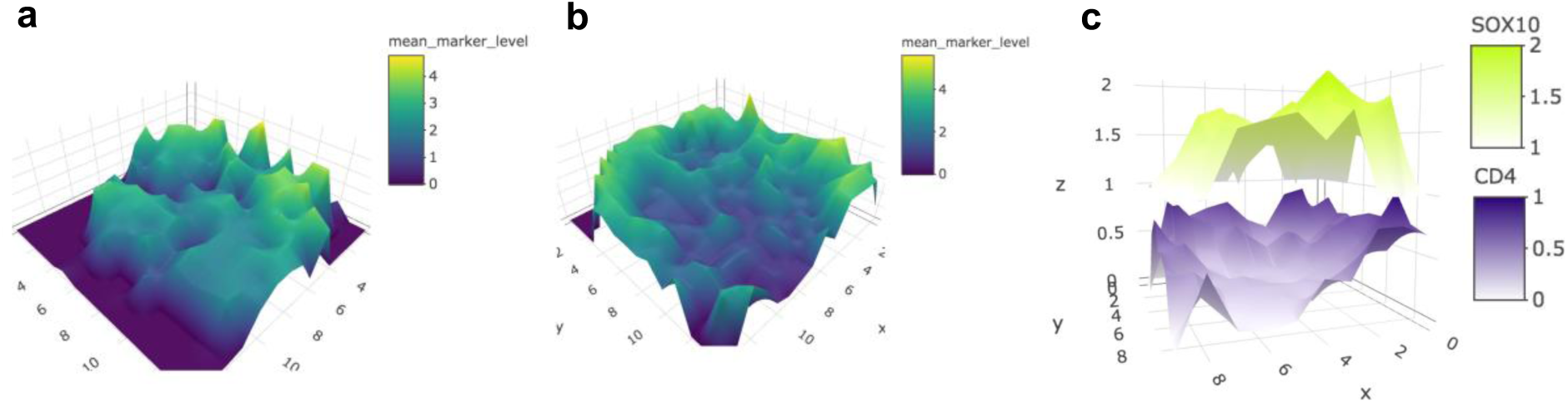
Surface plots of the SOX10 (a) and CD4 (b) markers in the melanoma tissue. Panel (c) shows how SOX10 and CD4 are mutually exclusive (valleys and mountains are opposite).

**Figure 6.**
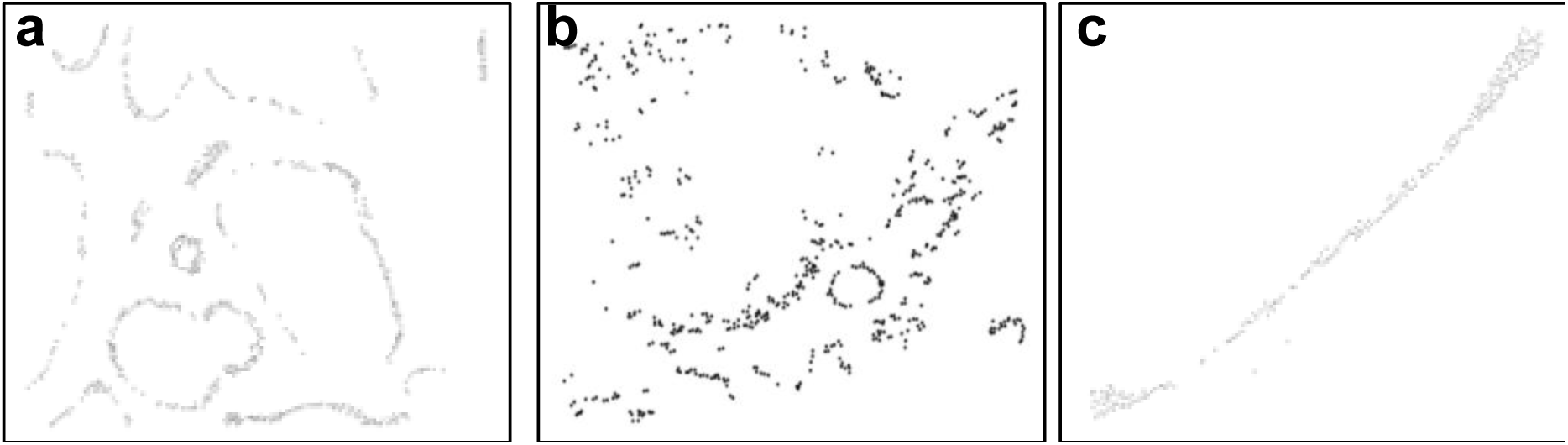
Bordering cells. (a-c) Bordering cells of the tissues in Figure 2b, 2c and 2d, respectively.

The calculation of the average minimum distance between all cell types revealed vastly different configurations between prostate cancer images (Figure 7). In the case of the image in Figure 2c, tumour cells were closely interacting with CD3+CD4+ and CD3+CD8+ cells (Figure 7a), suggesting higher levels of tumour-immune interaction. In contrast, the average minimum distance was much greater in Figure 7b (corresponding to the image in Figure 2d) suggesting immune exclusion. A similar analysis was performed looking at the distribution of minimum distances (Figure 8). Here, CD3+CD4+ closely interacting with the tumour were identified in the first image (Figure 8a), while this group of cells was not found in the latter (Figure 8b).

**Figure 7.**
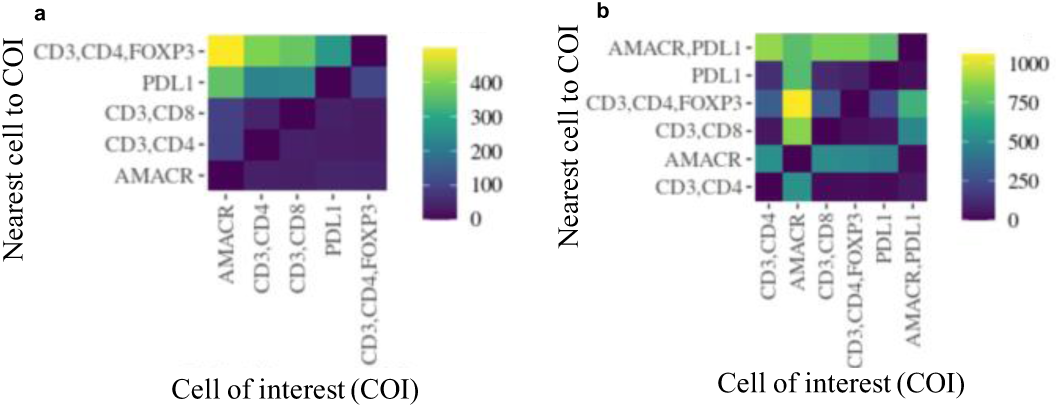
Average minimum cell distances between pairs of cell phenotypes. Panel (a) corresponds to Figure 2c, while panel (b) corresponds to Figure 2d. In (a) tumor (AMACR+) cells are closely interacting with CD3+CD4+ cells, whereas in (b) they are distant.

**Figure 8.**
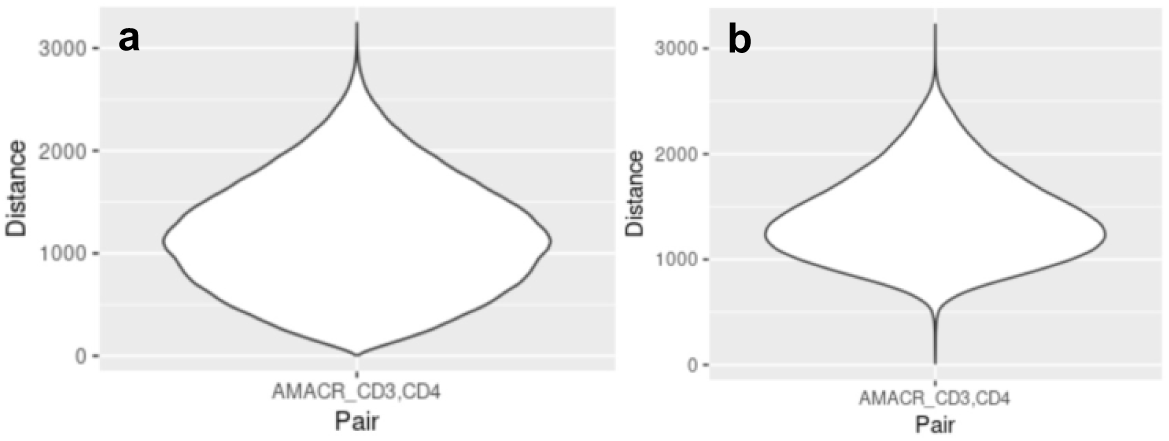
Average minimum cell distances between pairs of cell phenotypes. Panel (a) corresponds to Figure 2c, while panel (b) corresponds to Figure 2d. In (a) tumor (AMACR+) some cells are closely interacting with CD3+CD4+ cells, whereas in (b) most are distant.

We next investigated whether we could detect clusters of immune cells, and whether there was co-occurrence of immune and tumour cells in communities (Figure 9). Here, we define clusters as being composed of a particular cell phenotype or combination of phenotypes selected *a priori* by the user. In this example we selected to detect clusters of CD3+CD4+ and CD3+CD8+ cells in our prostate cancer images (Figure 9a-c). In contrast, communities refer to aggregates of cells identified solely by their location in the image, regardless of phenotype, as can be observed in (Figure 9d-f).

**Figure 9.**
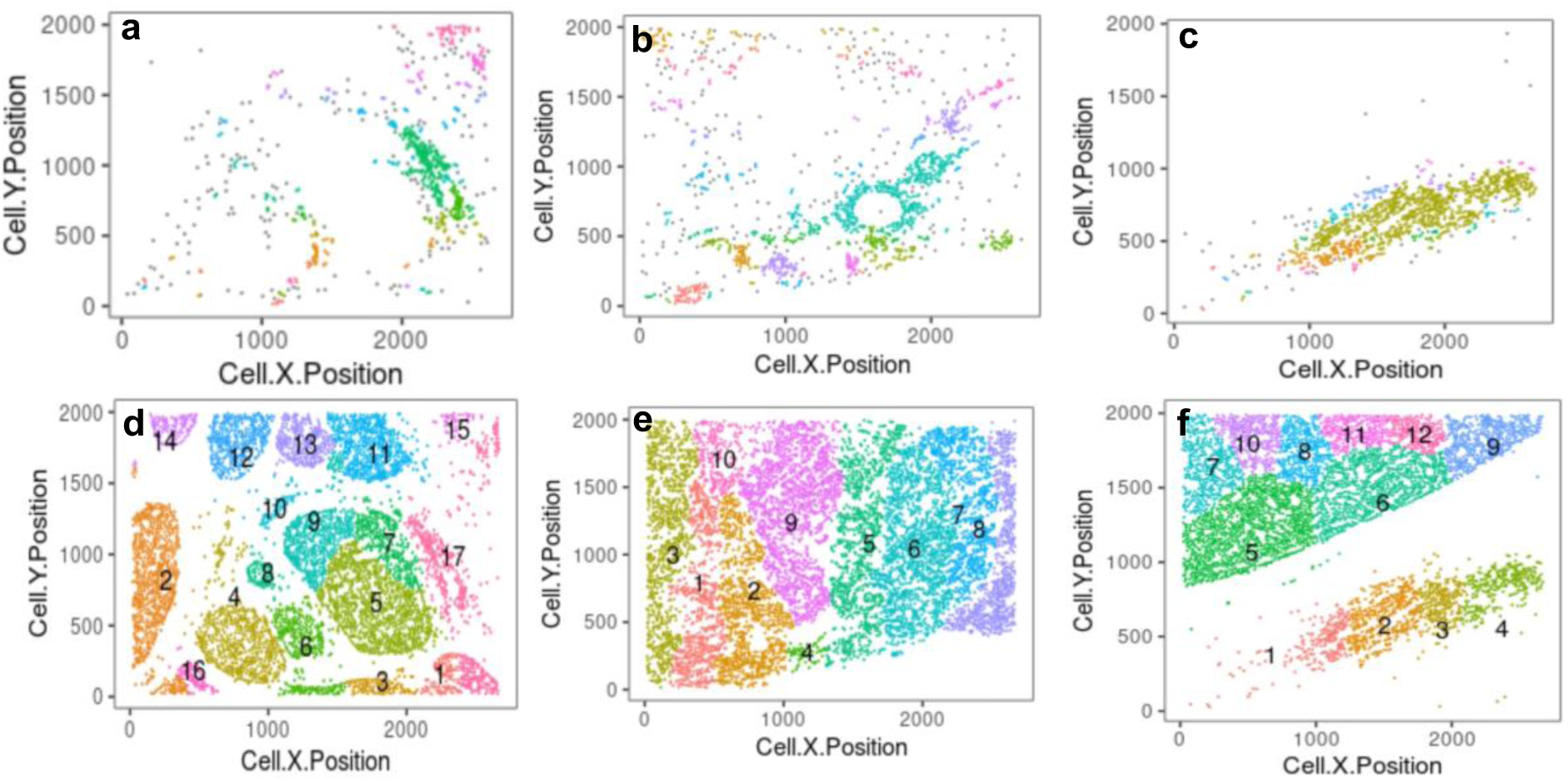
Cell clusters and communities in prostate cancer tissues. (a-c) Clusters of CD3+CD4+ and CD3+CD8+ cells. Each colour corresponds to a distinct cluster. Grey cells correspond to ‘free’, un-clustered cells. (d-f) Cell communities detected using all cells. Each colour corresponds to a community. Tissues correspond to Figures 2b-d.

Investigation into the composition of clusters and communities provided insights into how diverse cell types are aggregated in the tissues shown in Figure 2c and Figure 2d. While most clusters of the image in Figure 2c were composed of a mixture of CD3+CD4+ and CD3+CD8+ cells, a subset were dominated by either cell type (Figure 10a). All communities were composed of a mixture of tumour and immune cells, suggesting immune infiltration across the entire image (Figure 10b). Interestingly, PDL1+ and FOXP3+ cells (Tregs) were found to co-occur in communities with the highest levels of CD3+CD4+ cells (Figure 10c). In the case of the image of Figure 2d, while most clusters have similar proportions of CD3+CD4+ and CD3+CD8+ cells (Figure 10d), there was a clear distinction of tumour and immune communities, with no mixing (Figure 10e).

**Figure 10.**
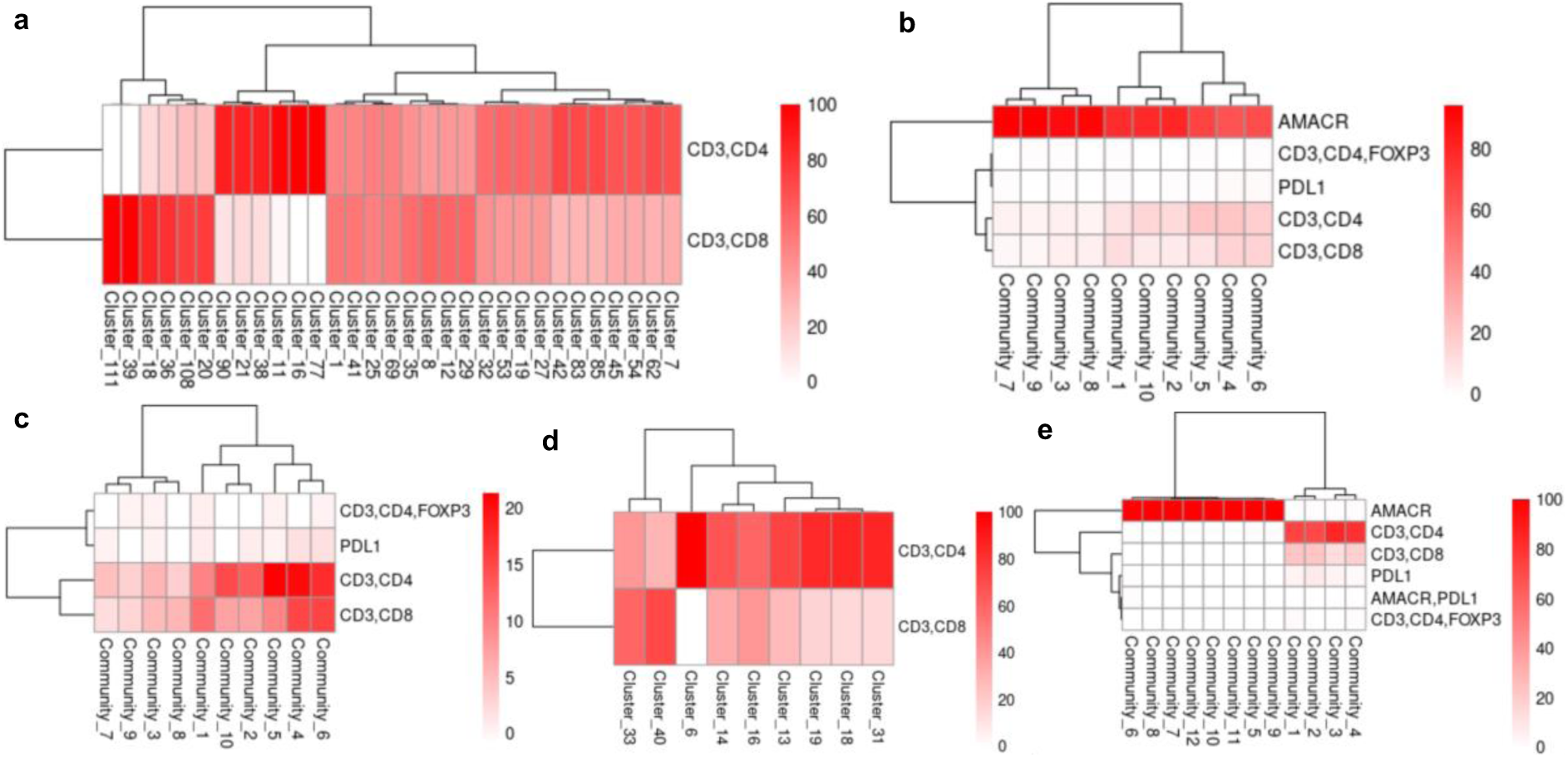
Cluster and community composition. (a) Cluster composition of tissue in Figure 2c. While most clusters are composed of a mixture of CD3+CD4+ and CD3+CD8+ cells, a subset are dominated by either. (b) Community composition of tissue in Figure 2c. All communities have a combination of tumour (AMACR+) and immune cells. (c) Same heatmap shown in panel (b), but without the AMACR marker. PDL1+ and FOXP3+ cells tend to appear in communities with high levels of CD3+CD4+ cells. (d) Cluster composition of tissue in Figure 2d. The vast majority of clusters include a mix of CD3+CD4+ and CD3+CD8+ cells. (e) Community composition of tissue in Figure 2d. There is a clear separation between communities with AMACR+ cells and those without.

These results are consistent with those obtained the mixing scores of these images. The mixing score CD4+ and tumour cells of the image of Figure 2c was 0.022, while for the image of Figure 2d it was 0.00095, suggesting higher levels of tumor-immune cell mixing in the former (Figures 10b and 10e).

To characterize the interaction of immune cells across our images, we performed gradient analysis with SPIAT. We investigated the intensity of the CD4 marker at different radii from from CD8+ cells. We found that the intensity of CD4 in cells surrounding CD8+ cells was low in the melanoma image, but that this intensity increased as the distance to CD8+ cells increased (Figure 11a). This suggested that CD4+ and CD8+ cells are likely forming their own separate clusters, which can be qualitatively perceived in Figure 2a. In contrast, for the 3 prostate cancer images, higher levels of CD8 were observed in cells closely interacting with CD4+ cells (Figure 11b-d) but these levels decreased at larger radii. This suggested a higher level of mixing of CD4+ and CD8+ cells in these images.

**Figure 11.**
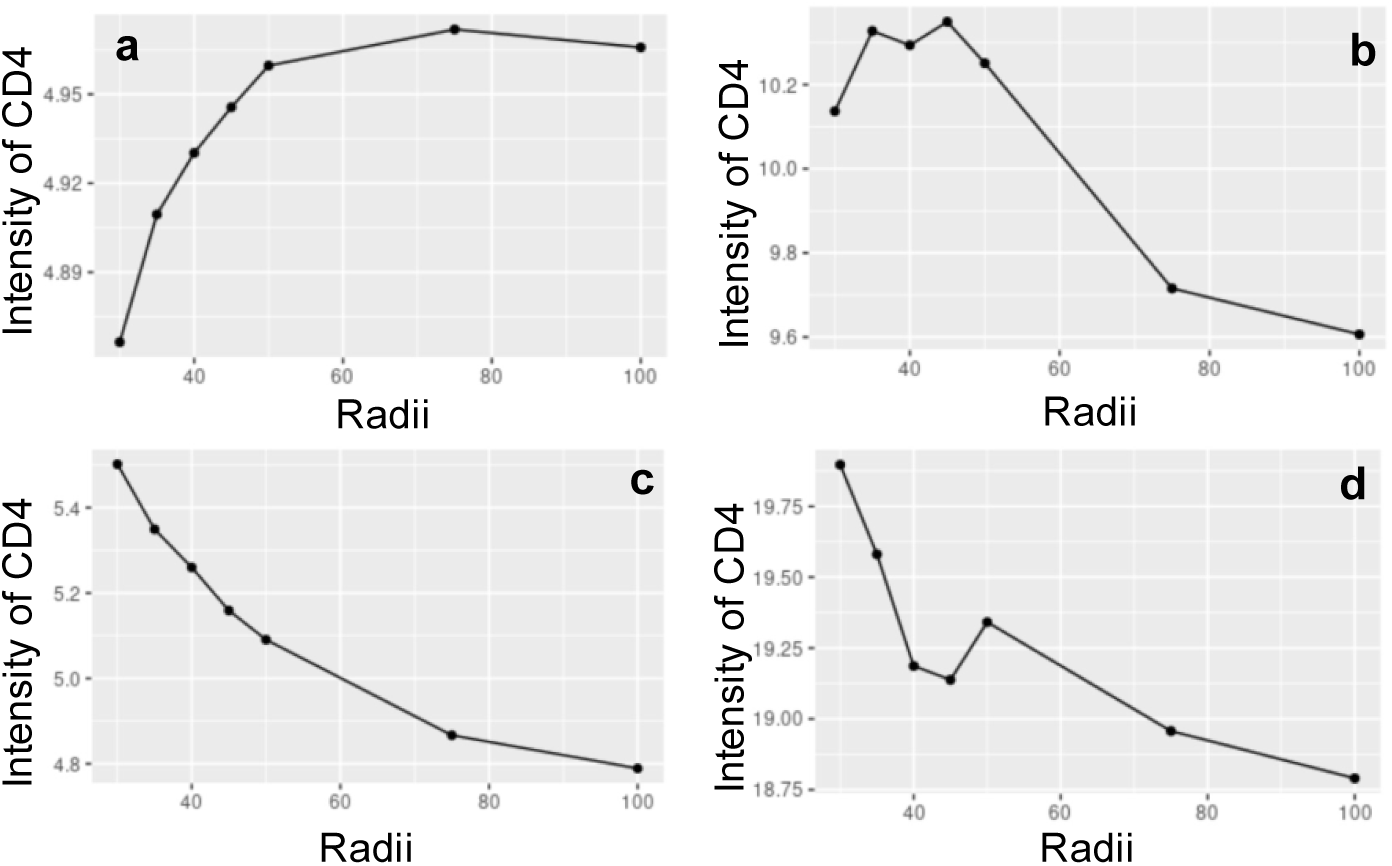
Average intensity of CD4 at various radii from CD8+ cells. (a-d) Panels correspond to those in Figure 2. In melanoma (a) the level of CD4 increases further from CD8+ cells due to segregation of CD4 and CD8 cells, whereas in the prostate cancer tissues (b-d) there is mixing of both cell types.

Finally, we tested the ability of SPIAT to automatically phenotype tumour cells based on marker intensities. We found that the spatial distribution of automatically phenotyped tumour cells resembled that with the phenotypes obtained using HALO and InForm (Figure 12). For the melanoma image, the number of true positive cells was 127,750 and true negative cells was 93,233, with only 3,198 false positives and no false negatives. For the prostate image, there were 5,441 true positive cells and 1,194 true negative cells, with only 79 false positives and 157 false negatives.

**Figure 12.**
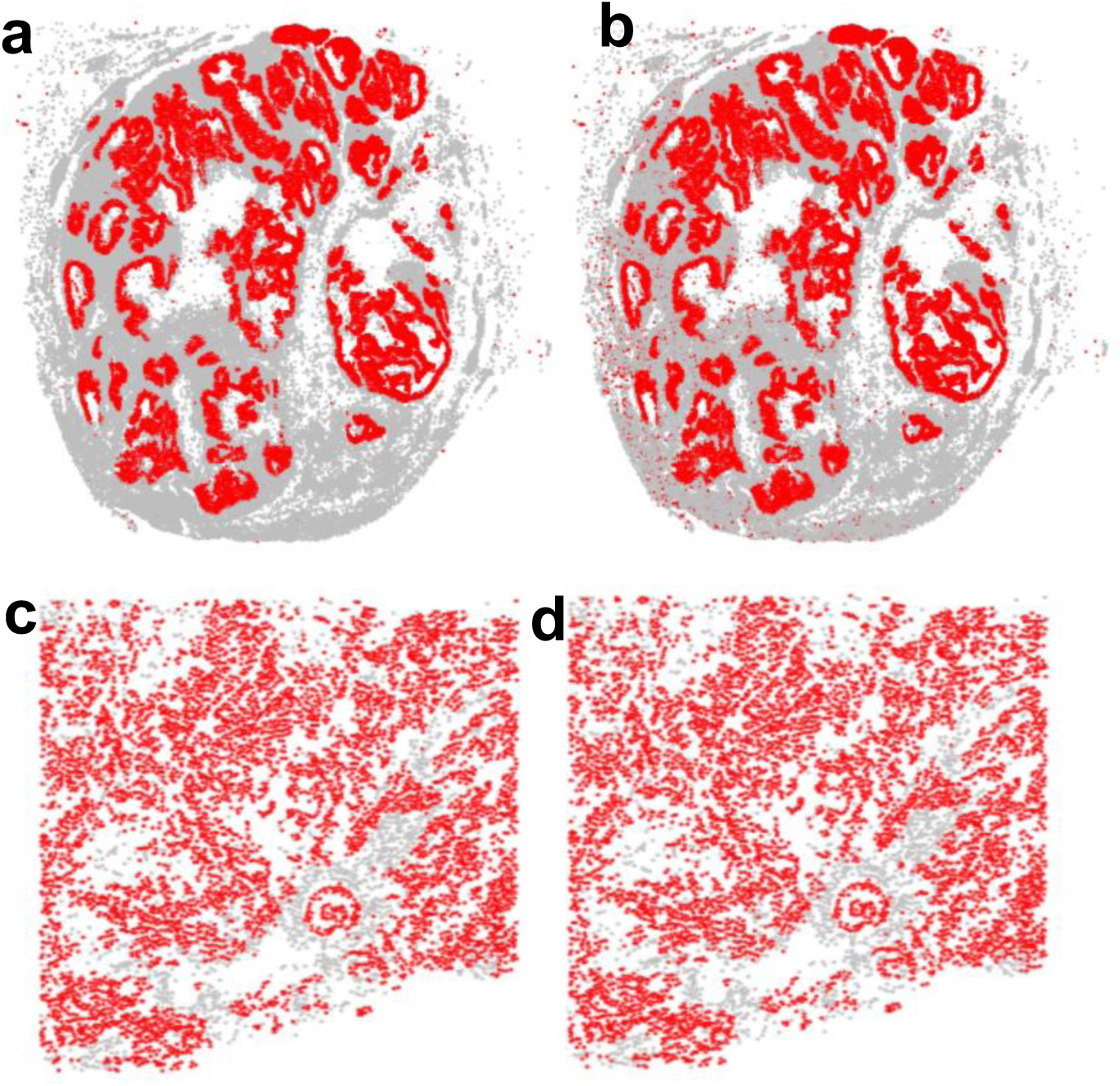
Prediction of tumour markers based on marker intensities. (a) SOX10+ cells as defined by HALO in melanoma tissue in Figure 2a. (b) Predicted SOX10+ cells using SPIAT. (c) AMACR+ cells in tissue of Figure 2c defined by InForm. (d) Predicted AMACR+ cells using SPIAT. Tumour cells are in red, other cells are in grey.

## Conclusion

SPIAT provides a broad range of tools for the spatial analysis of cells in tissues. While we have focused on the study of the tumour microenvironment, its application can be extended to other studies investigating the spatial distribution of cells in tissues. SPIAT continues to be in development, where we plan to further extend the number of metrics to characterize individual images, as well as include statistical tests to compare images.

## Acknowledgements

We would like to thank George Au-Yeung, Nineveth Rashoo and Richard Young for generously providing OPAL images for the development of this package, as well as Graeme Sissing and Steve Cavil for their support.

## Contributions

TY: Implemented algorithms; VO: Implemented algorithms; APa: Generated prostate cancer images; NK: Generated melanoma images; APi: Generated melanoma images; YH: Suggested metrics and algorithms; GB: Designed algorithms; SK: General supervision and advice on spatial patterns; PN: General supervision and advice on metrics; SS: General supervision and prostate cancer images; DG: General bioinformatics supervision; AT: Conceived project, designed and implemented algorithms, designed package, wrote manuscript.

## Notes

### Competing Interest Statement

The authors have declared no competing interest.

